# A novel highly potent inhibitor of TMPRSS2-like proteases blocks SARS-CoV-2 variants of concern and is broadly protective against infection and mortality in mice

**DOI:** 10.1101/2021.05.03.442520

**Authors:** Tirosh Shapira, I. Abrrey Monreal, Sébastien P. Dion, Mason Jager, Antoine Désilets, Andrea D. Olmstead, Thierry Vandal, David W. Buchholz, Brian Imbiakha, Guang Gao, Aaleigha Chin, William D. Rees, Theodore Steiner, Ivan Robert Nabi, Eric Marsault, Julie Sahler, Avery August, Gerlinde Van de Walle, Gary R. Whittaker, Pierre-Luc Boudreault, Hector C. Aguilar, Richard Leduc, François Jean

**Author notes:** **Corresponding Authors:** Hector C. Aguilar^2^, Richard Leduc^3^, and François Jean^1^. These two authors contributed equally (Tirosh Shapira, I. Abrrey Monreal).

## Abstract

The COVID-19 pandemic caused by the SARS-CoV-2 virus remains a global public health crisis. Although widespread vaccination campaigns are underway, their efficacy is reduced against emerging variants of concern (VOCs) ^1,2^. Development of host-directed therapeutics and prophylactics could limit such resistance and offer urgently needed protection against VOCs ^3,4^. Attractive pharmacological targets to impede viral entry include type-II transmembrane serine proteases (TTSPs), such as TMPRSS2, whose essential role in the virus lifecycle is responsible for the cleavage and priming of the viral spike protein ^5–7^. Here, we identify and characterize a small-molecule compound, N-0385, as the most potent inhibitor of TMPRSS2 reported to date. N-0385 exhibited low nanomolar potency and a selectivity index of >10^6^ at inhibiting SARS-CoV-2 infection in human lung cells and in donor-derived colonoids ^8^. Importantly, N-0385 acted as a broad-spectrum coronavirus inhibitor of two SARS-CoV-2 VOCs, B.1.1.7 and B.1.351. Strikingly, single daily intranasal administration of N-0385 early in infection significantly improved weight loss and clinical outcomes, and yielded 100% survival in the severe K18-human ACE2 transgenic mouse model of SARS-CoV-2 disease. This demonstrates that TTSP-mediated proteolytic maturation of spike is critical for SARS-CoV-2 infection *in vivo* and suggests that N-0385 provides a novel effective early treatment option against COVID-19 and emerging SARS-CoV-2 VOCs.

## Introduction

In December 2019, the first cases of coronavirus disease 2019 (COVID-19) emerged in Wuhan, Hubei Province, China, and were rapidly attributed to the etiology of a novel β-coronavirus, severe acute respiratory syndrome coronavirus 2 (SARS-CoV-2) ^9^. As of May 3, 2021, more than 153 million SARS-CoV-2 infections and over 3.2 million deaths have been reported ^10^. The approval and widespread distribution of several highly effective vaccines, along with other public health measures, now allows the possibility of controlling the COVID-19 pandemic; however, novel genetic variants of SARS-CoV-2 are emerging and spreading at alarming speed ^11^. Importantly, vaccine effectiveness may be reduced against a number of these variants, termed *variants of concern* (VOCs) ^2,12,13^.

Discovery of novel classes of antiviral compounds including both direct-acting (DAA) and host-directed (HDA) antivirals and intensive *in cellulo* and *in vivo* studies of their antiviral profiles as mono- or combination therapies against emerging SARS-CoV-2 VOCs are critical for developing preventive and therapeutic strategies to combat COVID-19 ^6,14,15^. Remdesivir is the only antiviral currently approved for clinical use against SARS-CoV-2 ^16^. Remdesivir is a DAA targeting the viral RNA-dependent RNA polymerase that catalyzes the synthesis of viral RNA ^17^. Remdesivir is currently administered intravenously to hospitalized patients with COVID-19 ^16^. Another DAA, PF-07321332 is being developed as an oral clinical candidate. It targets the coronavirus’s main protease (M^pro^, also called 3CL^pro^), an essential protease involved in processing viral replicase polyproteins ^18^. Alternatively, host-directed antivirals (HDAs) (also called indirect-acting antivirals) are under investigation and may offer a complement to DAAs. HDAs have reduced potential for resistance by emerging SARS-CoV-2 VOCs since unlike viral genes, host genes possess a low propensity to mutate compared to viral genes ^5,6^. Camostat mesylate (Cm), for example, is a repositioned clinical candidate for treating COVID-19 that is targeted at human type-II transmembrane serine proteases (TTSPs) such as TMPRSS2 ^19^. Cm is a broad-spectrum serine protease inhibitor used to treat pancreatitis and has demonstrated activity against TTSPs, host proteases under active investigation as therapeutic targets for COVID-19 ^4,5^. The transgenic human SARS-CoV-2 receptor (angiotensin-converting enzyme 2 [hACE2]) under a cytokeratin 18 promoter (K18); K18-hACE2) mouse model offers a stringent system for testing the efficacy of DAAs and HDAs against severe disease and mortality following SARS-CoV-2 infection ^20^. To date, very few studies have tested antiviral efficacy in this animal model with only one DAA, a viral 3CL^pro^ inhibitor, reported as protecting against lethal SARS-CoV-2 infection in this model ^21–23^.

To date, accumulating evidence has demonstrated SARS-CoV-2 dependence on host pathways including the viral hijacking of TMPRSS2-related proteases for viral entry, suggesting that TTSPs are attractive therapeutic targets to prevent SARS-CoV-2 infection ^5^. The SARS-CoV-2 lifecycle begins with attachment and entry into respiratory epithelium via the ACE2 receptor ^4,9^. This is mediated by the major viral surface glycoprotein, spike (S), which must undergo two sequential proteolytic cleavages by host proteases before it can mediate fusion of the virus with host cell membranes, a requirement for subsequent viral replication ^3,24,25^. The first spike cleavage occurs at the S1/S2 site, releasing S1 and S2 subunits that remain non-covalently linked, an event potentially mediated by host furin-like proteases ^24,25^. The second cleavage occurs at the S2’ site, immediately adjacent to the fusion peptide. This cleavage, which triggers the fusion event, is mediated by host TTSPs, such as TMPRSS2 and TMPRSS13, which cleave after specific single arginine or lysine residues (Arg/Lys↓) ^4,7,26^.

Here, we report on the design and testing of novel small-molecule peptidomimetics for their inhibitory activity against TMPRSS2 and related TTSPs. We then investigated their broad-spectrum antiviral activity against SARS-CoV-2 and two VOCs (B.1.1.7 and B.1.351) in human cells. Last, we tested our lead highly potent antiviral, N-0385, against SARS-CoV-2-induced morbidity and mortality in K18-hACE2 mice, a model of severe COVID-19.

## Results

### Small-molecule peptidomimetics with ketobenzothiazole warheads are potent inhibitors of TMPRSS2

We previously designed first-generation peptidomimetic tetrapeptide compounds having ketobenzothiazole warheads, which demonstrated inhibitory activity against a host TTSP, matriptase ^27,28^. These compounds act as slow tight-binding inhibitors *in vitro* but their potency in cellular systems was modest against influenza A virus ^28^. To improve their stability and potency, we modified their N-terminus either by capping or through synthesis of desamino moieties (**Figure 1A** and **Figure S1**) ^29^. When we measured the stability of desamino compounds, we found that they had drastically increased half-lives compared to their corresponding amine analogs (48 hr versus 2 hr, respectively, in human lung epithelial Calu-3 cells) (data not shown). Moreover, these compounds exhibited low nanomolar efficacies when tested in H1N1 models of influenza A virus infection ^28,30^.

**Figure 1.**
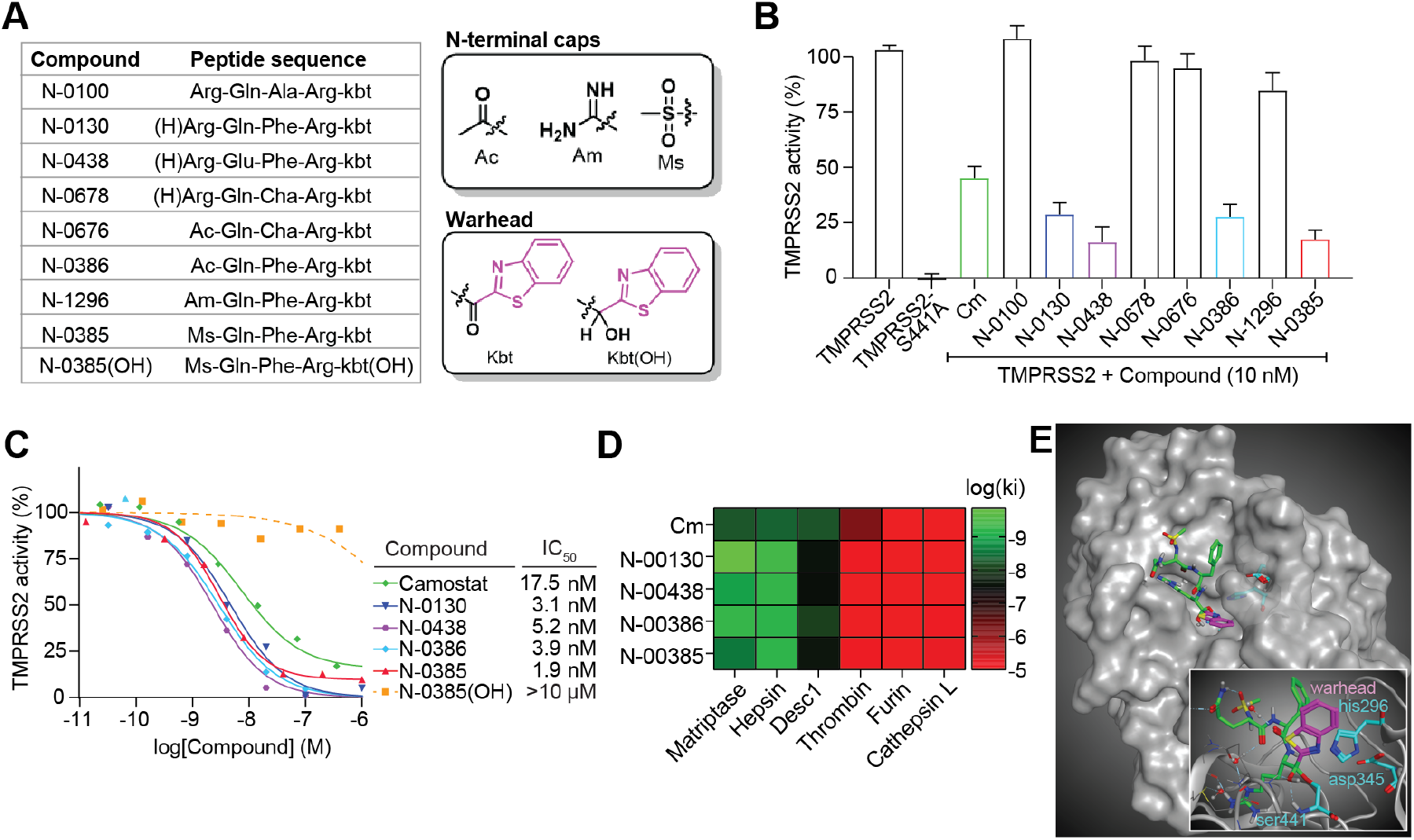
Ketobenzothiazole-based small-molecule peptidomimetics are potent TMPRSS2 inhibitors. **(A)** List of the peptidomimetic compounds used in this study along with their respective sequences. The structures of N-terminal caps, the ketobenzothiazole warhead, and the alcohol ketobenzothiazole are shown on the right. **(B)** Vero E6 cells were transfected with either an empty vector (mock), TMPRSS2 wild type (WT), or the inactive mutant TMPRSS2-S441A for 24 hr. Indicated compounds (10 nM) were added concomitantly with a fluorogenic substrate on cells for an additional 24 hr before fluorescence reading. Relative TMPRSS2 activity was measured using the mock-subtracted fluorescence and reported as the percentage of residual activity relative to the saline-treated cells (0.01% DMSO). n = 3. **(C)** Dose-response curves were generated for the indicated compounds (n ≥ 3) using the assay described in (A) and IC_50_ values were determined using nonlinear regression analysis. Representative IC_50_ curves are shown. **(D)** Specificity of selected compounds toward other serine proteases are shown. Data are represented as log (K_i_), n ≥ 3 and represented as a heat map. **(E)** Large: Docking of N-0385 (green, warhead in purple) in the binding pocket of TMPRSS2 (homology model). Residues of the catalytic triad are shown in cyan. Small: Interaction of N-0385 with TMPRSS2 residues. N-0385 forms a covalent bond with catalytic triad residue Ser441. (H)Arg = desamino arginine, Ac = acetyl, Am = amidinyl, Ms = mesyl, kbt = ketobenzothiazol, Cm = camostat mesylate, Cha = cyclohexylalanine

Expanding on that work here, we developed a small library of peptidomimetic compounds (**Figure 1A** and **Figure S1**) to screen for inhibition of TMPRSS2 proteolytic activity, as this TTSP is a crucial host protease involved in cleaving the SARS-CoV-2 spike and priming the virus for cell entry ^4^. We included in this screen our first-generation tetrapeptide, N-0100 ^28^, which lacks an N-terminal stabilizing group, along with three desamino tetrapeptide analogs. We also tested four tripeptides containing different N-terminal capping groups.

To evaluate the efficacies of these compounds, we set up a cellular assay to measure TMPRSS2-dependant pericellular inhibition of proteolytic activity. We expressed the full-length, wild type TMPRSS2 or an inactive form of the protease in which the serine residue of the catalytic triad was replaced by alanine (TMPRSS2-S441A) in Vero E6 cells. Twenty-four hr after transfection, the media was replaced for an additional 24 hr with serum-free media containing vehicle or compound in the presence of a TMPRSS2-preferred fluorogenic substrate ^31^ (**Figure 1B**). TMPRSS2-transfected cells treated with vehicle exhibited a 5-fold increase in fluorescent reporter activity compared to mock transfected cells, while TMPRSS2-S441A-expressing cells had no activity over background.

The peptidomimetics were tested for inhibitory activity against TMPRSS2 at 10 nM (**Figure 1B**). Camostat mesylate (Cm), which has previously been shown to be active against TMPRSS2 ^32^, reduced substrate proteolysis by 56% compared to untreated TMPRSS2-expressing cells. The first-generation peptidomimetic, N-0100, did not inhibit TMPRSS2 activity under these conditions. However, the more stable tetrapeptides with N-terminus desamino moieties, N-0130 and N-0438, had increased inhibitory activity of 72% and 84%, respectively. N-0678 (substituting P2 Phe for the synthetic amino acid Cha) only inhibited TMPRSS2 activity by 5%. N-0676 (a tripeptide with an N-terminal Ac cap and P2 Cha) also weakly inhibited TMPRSS2 activity by 8%. N-0386 (restoring Phe in P2) resulted in more potent inhibition of 73%. N-1296 (replacing Ac with Am) had reduced potency of 16%, while N-0385 (replacing Am with Ms) resulted in a highly potent inhibition of 83%. Importantly, several peptidomimetic compounds were more efficient than Cm at reducing TMPRSS2 activity (**Figure 1B**).

We then investigated the dose response of the four most promising peptidomimetics (N-0130, N-0385, N-0386, and N-0438). The half-maximal inhibitory concentration (IC_50_) of Cm was 17.5 ± 18.8 nM, while the IC_50_ for N-0130 was 3.1 ± 1.5 nM; for N-0438 it was 5.2 ± 5.4 nM; for N-0386 it was 3.9 ± 4.4 nM; and for N-0385 it was 1.9 ± 1.4 nM (**Figure 1C** and **Table S1**). Importantly, none of the compounds affected Vero E6 cellular viability when used at 10 µM (**Figure S2**). To confirm the contribution of the ketobenzothiazole warhead to the molecule’s inhibitory activity, the ketone functional group of N-0385 was replaced with an alcohol group to generate N-0385(OH) (**Figure 1A**), which we predict no longer traps the target protease. No significant reduction in TMPRSS2 activity was detected when cells were treated with up to 10 µM of N-0385(OH) (**Figure 1C**), suggesting that the integrity of the ketobenzothiazole group is required to achieve potency.

Next, we sought to determine the selectivity profile of these inhibitors by measuring the dissociation constant *K*_i_ on selected recombinant serine proteases, including three members of the TTSP family (matriptase, hepsin, DESC1) as well as furin, thrombin, and cathepsin L. All four peptidomimetic compounds we tested behaved as low nanomolar inhibitors for the TTSPs, but they were inactive or showed only weak inhibition against the other proteases (**Figure 1D** and **Table S2**). Cm displayed a similar selectivity profile to the peptidomimetics tested, except that it demonstrated moderate inhibition of thrombin (*K*_i_ = 621 nM) in line with its broader spectrum properties. Overall, TTSP-targeting peptidomimetics harbouring a ketobenzothiazole warhead inhibit TMPRSS2-dependent pericellular activity in a cellular assay and preferentially inhibit other members of the TTSP family.

To understand the mode of binding and the main interactions of our inhibitors and how these compounds achieve their high inhibitory potential, we built a homology model of TMPRSS2 using the crystal structure of matriptase (PDB: 6N4T). Alignment of the catalytic domains demonstrated 41% and 60% identity and sequence similarity, respectively, making it a reliable model, especially near the conserved binding site. Docking of N-0385 was modeled to this structure (**Figure 1E**). As predicted and recently published ^33^, the catalytic triad Ser441 (catalytic triad: Ser441, His296, and Asp345; **Figure 1E** inlet) forms a covalent bond with the warhead ketone, thus leading to a tight-binding mode of inhibition.

Several key interactions can be observed in the binding pocket. As in all TTSP inhibitors possessing a guanidine group on the side-chain, a strong hydrogen bond network stabilizes this pharmacophore deep within the binding pocket (**Figure 1E**). This includes Asp435 and Gly464 as well as Gln438 via a water molecule. Gln438 is also involved in another hydrogen bond of this same water molecule to the inhibitor’s glutamine ketone. This ketone also acts as a hydrogen bond acceptor with Gly462. The N-terminal mesylate forms two hydrogen bonds, one intramolecular with the terminal amide of N-0385 and another with Gly462. Finally, the oxygen of the newly formed hemiacetal is stabilized by two hydrogen bond donors from the Gly439 and Ser441 amines. A portion of the ketobenzothiazole warhead and the aromatic ring from the phenylalanine are exposed to the solvents, which could allow us to further optimize the design of this second-generation inhibitor leading to an improved pharmacokinetic profile.

### Small-molecule peptidomimetics with ketobenzothiazole warheads are potent inhibitors of SARS-CoV-2 entry in a lung epithelial cell line and in donor-derived human colonoids

The peptidomimetic compounds we screened against TMPRSS2 were subsequently tested for their efficacy at preventing SARS-CoV-2 infection. Calu-3 cells were pretreated with 100 nM of the compounds for three hr prior to infection. Cells were fixed and immunofluorescently stained for dsRNA, a marker of viral replication ^34^, and for the viral nucleocapsid, a marker of viral entry and translation ^35^ (**Figure S3**). Fluorescent high-content imaging and relative quantification of virally infected cells demonstrated consistent inhibitory profiles across dsRNA and nucleocapsid staining, which mirrored the inhibitory profile observed in the TMPRSS2 proteolytic activity assay (**Figure 2A** versus **Figure 1B**). Cm, which interferes with SARS-CoV-2 infection ^4^, reduced infection by >83% compared to non-treated cells. N-0100, which lacks an N-terminal stabilizing moiety, reduced infection by <25%. The tetrapeptides N-0130 and N-0438, which have N-terminus desamino moieties, had greatly increased antiviral activity of >93% and >88%, respectively. N-0678 (substituting P2 Phe for the synthetic amino acid Cha) inhibited SARS-CoV-2 by <23%. N-0676 (tripeptide with an N-terminal Ac cap and P2 Cha) had only moderate inhibitory activity of <53%. N-0386 (restoring Phe in P2) resulted in a highly potent SARS-CoV-2 inhibition of >99%. N-1296 (replacing Ac with Am) reduced the antiviral potency to <44%, while N-0385 (capping with Ms) restored antiviral activity to >99%. Last, N-0385(OH) (OH replacing the functional group of the warhead), demonstrated a <23% inhibition of SARS-CoV-2. Thus, TMPRSS2-inhibiting peptidomimetics are also inhibitors of SARS-CoV-2 replication and translation in Calu-3 cells; the stabilizing N-terminal caps and the ketobenzothiazole warhead are likely essential for compound stability and antiviral potency.

**Figure 2.**
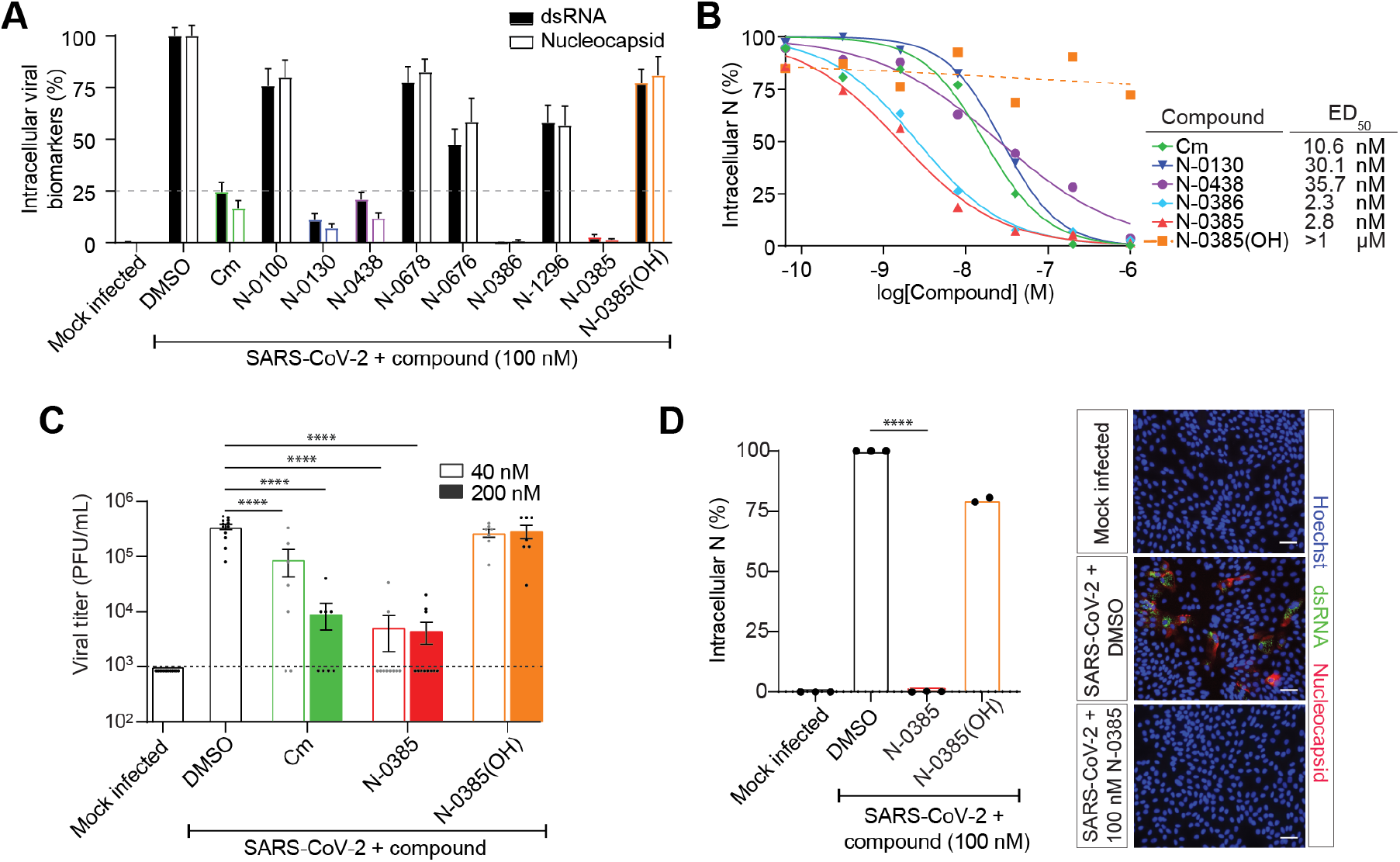
Small-molecule peptidomimetics active against TMPRSS2 are potent low nanomolar inhibitors of SARS-CoV-2 in a human lung epithelial cell line and in human colonoids. **(A)** Calu-3 cells were pretreated with 100 nM of the indicated compounds followed by SARS-CoV-2 infection (MOI = 2). Intracellular infection levels were evaluated by high-content screening of cell nuclei, dsRNA, and nucleocapsid and then quantified relative to DMSO-treated cells. (nucleocapsid, n = 3; dsRNA, n = 2); significant nucleocapsid comparisons: **** (modified p <0.0001) Cm, N-0130, N-0438, N-0676, N-0386, N-1296, and N-0385; significant dsRNA comparisons: * (modified p <0.05) N-1296, *** (modified p <0.0005) N-0676, **** Cm, N-0130, N-0438, N-0386, and N-0385. **(B)** Dose response curves were generated for the lead antiviral peptidomimetic compounds in Calu-3 cells using nucleocapsid staining of compound pretreated and infected cells (n: Cm = 5, N-0130 = 5, N-0438 = 3, N-0386 = 4, N-0385 = 8, N-0385(OH) = 5). **** (modified p <0.0001). **(C)** Plaque assays were performed using two of the experimental conditions evaluated in the dose response analysis (40 nM and 200 nM) to determine the viral titers (amount of infectious virus) produced in cells pretreated with the indicated compounds prior to infection (n = 3); dotted line represents limit of detection. **** (modified p <0.0001). **(D)** Colonoids were pretreated with 100 nM of the indicated compounds and infected with SARS-CoV-2 (MOI ≈ 1). The impact on intracellular infection levels was determined by relative quantification of nucleocapsid staining. Representative fluorescent images of colonoids subjected to the indicated treatments are shown (Hoechst in blue, nucleocapsid in red, and dsRNA in green). Scale bars 50 µm. (N-0385, n = 3; N-038(OH), n = 2); **** modified p <0.0001. One-way ANOVA with Bonferroni correction was used to determine significance in (A), (C), and (E). Error bars represent standard error of the mean. Cm = camostat mesylate, N = nucleocapsid, PFU = plaque-forming units, MOI = multiplicity of infection

Compounds significantly inhibiting SARS-CoV-2 (>75%) in the antiviral screen were further validated and characterized using a dose response analysis in Calu-3 cells (**Figure S3**). The half-maximal effective dose (ED_50_) of Cm was 10.6 ± 8.4 nM, while the ED_50_ for N-0130 was 30.1 ± 30.1 nM; for N-0438 it was 35.7 ± 24.5 nM; for N-0386 it was 2.3 ± 1.7 nM; and for N-0385 it was 2.8 ± 1.4 nM (**Figure 2B**). An ED_50_ value could not be determined for N-0385(OH) as significant inhibition was not observed at concentrations up to 1 µM (**Figure 2B)**. These compounds did not exhibit any toxicity; all four compounds had half-maximal cytotoxic concentration (CC_50_) values of >1 mM in Calu-3 cells (**Table S1**). Thus, the selectivity index (SI) for these compounds (N-0130, N-0438, N-0386, and N-0385) was between 8.97 X10^4^ and 2.75 X10^6^ (**Table S1**). Overall, these results confirm that two newly discovered TTSP-targeted peptidomimetic compounds (N-0386 and N-0385) are extremely potent low nanomolar inhibitors of SARS-CoV-2 infection in human lung epithelial cells.

We next examined the impact of Cm, N-0385, and N-0385(OH) on the extracellular release of SARS-CoV-2 infectious virions from Calu-3 cells. Two effective doses (40 nM and 200 nM) from the ED_50_ curve-fitting (**Figure 2B)** were selected for plaque assays. The cell supernatant from Cm-treated and infected cells demonstrated a 1-log reduction in the presence of 40 nM compared to the DMSO-treated infected control and a 2-log reduction with 200 nM Cm (**Figure 2C**). In comparison, both 40 nM and 200 nM treatments with N-0385 reduced viral titers by >2.5-log. Consistent with previous results, N-0385(OH) did not exhibit a significant reduction in SARS-CoV-2 plaques at 40 nM or 200 nM. These results confirm that N-0385, which targets TMPRSS2, is a potent inhibitor of SARS-CoV-2 infectivity in Calu-3 cells and that the ketobenzothiazole warhead is required for N-0385 antiviral potency.

Although Calu-3 cells represent a scalable and clinically relevant system of antiviral screening for SARS-CoV-2 inhibitors, they are an immortalized cell line ^36^. To evaluate the effectiveness of N-0385 in a primary human cell-based model, we explored SARS-CoV-2 infection in donor-derived human colonoids ^37,38^. SARS-CoV-2 initially causes a respiratory infection but many infected individuals also experience gastrointestinal symptoms frequently linked with increased disease duration and severity ^39,40^. We first investigated the susceptibility of colonoid monolayers ^37^ to SARS-CoV-2 infection. Consistent with previous work, the colonoids were susceptible to infection as evidenced by dsRNA and nucleocapsid staining (**Figure 2D** and **Figure S4**) ^39,40^.

N-0385 and N-0385(OH) were then tested for their efficacy at preventing SARS-CoV-2 infection in colonoids. The colonoids were pretreated with 100 nM of the compounds for 3 hr prior to 3-day infection with SARS-CoV-2. Under these conditions, N-0385-pretreated colonoids had undetectable infection compared to DMSO-treated colonoids (>99% inhibition) (**Figure 2D)**. In contrast, N-0385(OH) did not significantly reduce SARS-CoV-2 infection in this system (<20% inhibition) (**Figure 2D)**. These results align with observations in Calu-3 cells and confirm the nanomolar potency of N-0385 against SARS-CoV-2 in primary human cells.

### N-0385 is a nanomolar, broad-spectrum coronavirus inhibitor of SARS-CoV-2 VOCs including B.1.1.7 and B.1.351

To our knowledge, mutations in the TMPRSS2 cleavage site have not been identified in SARS-CoV-2 variants, suggesting that that N-0385 should retain high potency against SARS-CoV-2 VOCs ^12^. First, we confirmed infectivity of two VOCs in Calu-3 cells: B.1.1.7 (originally identified in the United Kingdom) and B.1.351 (first identified in South Africa). Confocal imaging of infected cells confirmed infectivity of these variants as demonstrated by nucleocapsid and dsRNA staining (**Figure 3A**). We then evaluated the efficacy of N-0385 for preventing SARS-CoV-2 VOC infection in Calu-3 cells. The ED_50_ of N-0385 against the VIDO-01 isolate was 5.2 nM under these conditions while the ED_50_ against B.1.1.7 was 3.4 nM and against B.1.351 it was 13.1 nM (**Figure 3B**). Statistical analysis confirmed that compared to the VIDO-01 isolate, there was no difference in effectiveness of N-0385 against the B.1.1.7 variant; however, there was a significant difference of N-0385 against the B.1.351 variant in Calu-3 cells. In both cases, N-0385 retained low nanomolar potency against both VOCs. This underlines the potential of N-0385 to act as a broad spectrum, host-directed antiviral against emerging SARS-CoV-2 VOCs.

**Figure 3.**
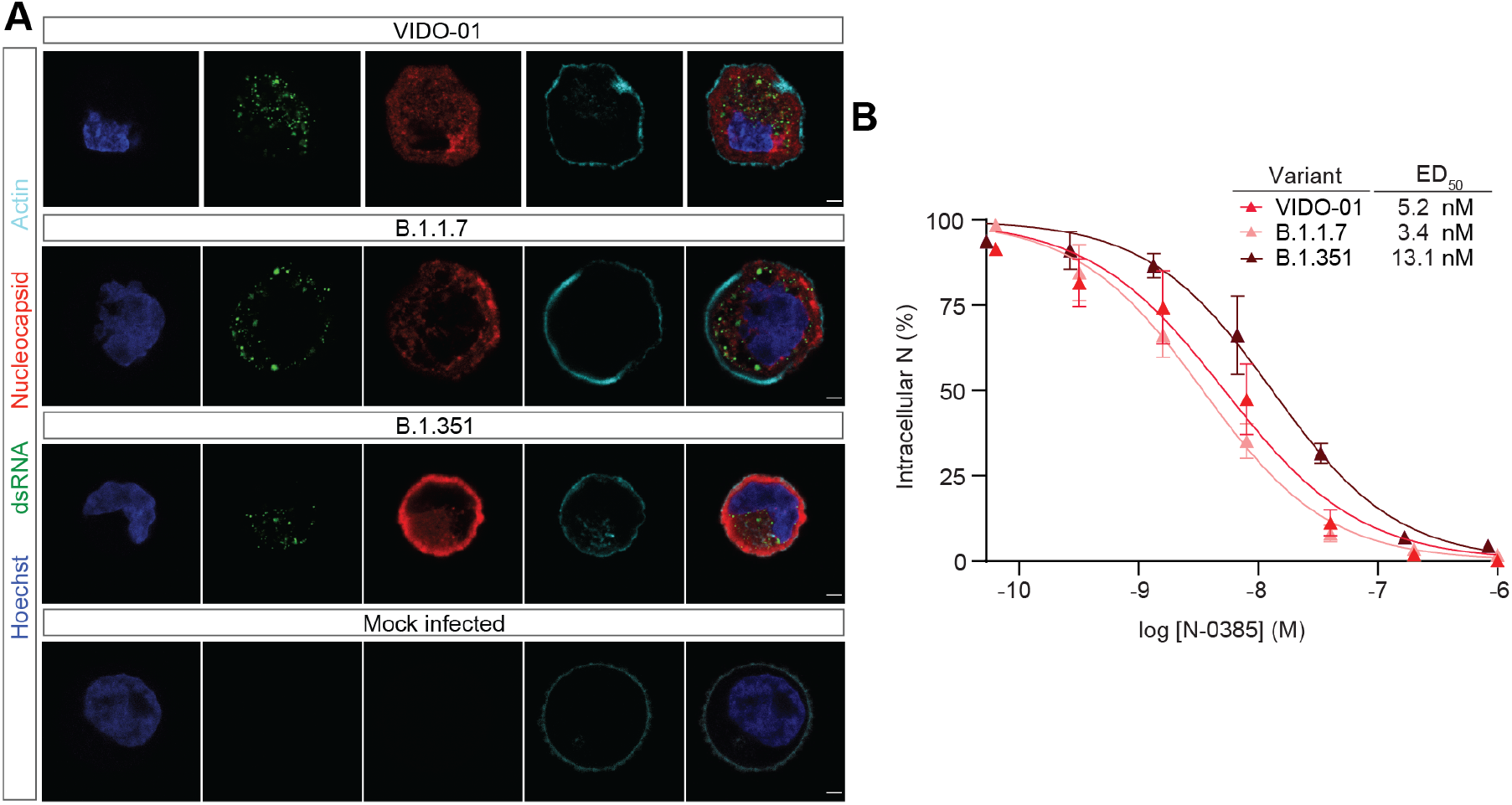
N-0385 is a nanomolar broad-spectrum coronavirus inhibitor of SARS-CoV-2 variants of concern (VOCs). **(A)** Representative fluorescent images of SARS-CoV-2-infected Calu-3 cells. Calu-3 cells infected with the indicated SARS-CoV-2 variants and mock infected are shown. Scale bar: 5 µm. Hoechst is shown in blue, nucleocapsid in red, dsRNA in green, and actin in cyan. Images captured with a Leica TCS SP8 3× STED microscope. **(B)** Dose response curves were generated for N-0385 in Calu-3 cells using nucleocapsid staining of N-0385 pretreated cells infected with the indicated VOCs (n = 3). The significant difference between the dose response curves for B.1.1.7 and B.1.351 compared to the VIDO-01 isolate was assessed using repeated measures ANOVA with concentration as a random factor. Tukey’s post hoc analysis was used to test pair-wise comparisons. There was a significant main effect of the variant on the dose response curves (F = 7.0708, p = 0.0099). Post hoc analysis showed that the dose response curve for the B.1.351 variant, but not the B.1.1.7 variant, was significantly different from the VIDO-01 (p=0.0301) isolate.

### N-0385 prevents SARS-CoV-2-induced morbidity and mortality in a mouse model of infection

After establishing the efficacy of N-0385 *in vitro* and *in cellulo*, we tested whether intranasal administration would be protective *in vivo*, using K18-hACE2 mice expressing the human ACE2 receptor driven by a keratin promoter ^41,42^, an established mouse model of severe SARS-CoV-2 disease ^43^. K18-hACE2 mice express the human ACE2 receptor driven by a keratin promoter ^41,42^. In the first experiment, ten mice per group (five female and five male) were administered a single daily intranasal dose of 7.2 mg/kg N-0385, N-0385(OH), or a vehicle control (0.9% saline) for eight days from day -1 to day 6 relative to infection. The mice were challenged on day 0 with 1 x 10^3^ PFU/mouse of SARS-CoV-2 (**Figure 4A**). Weight loss and survival data indicate that mice receiving N-0385 exhibited greatly reduced morbidity and mortality compared to untreated mice (**Figure 4B-E**). N-0385-treated mice exhibited lower weight loss (average 3%) and greater survival (70%) in stark contrast with N-0385(OH)- or saline-treated mice, which had significantly greater weight loss (average ∼14% for both) and poor survival (10% and 0%, respectively).

**Figure 4.**
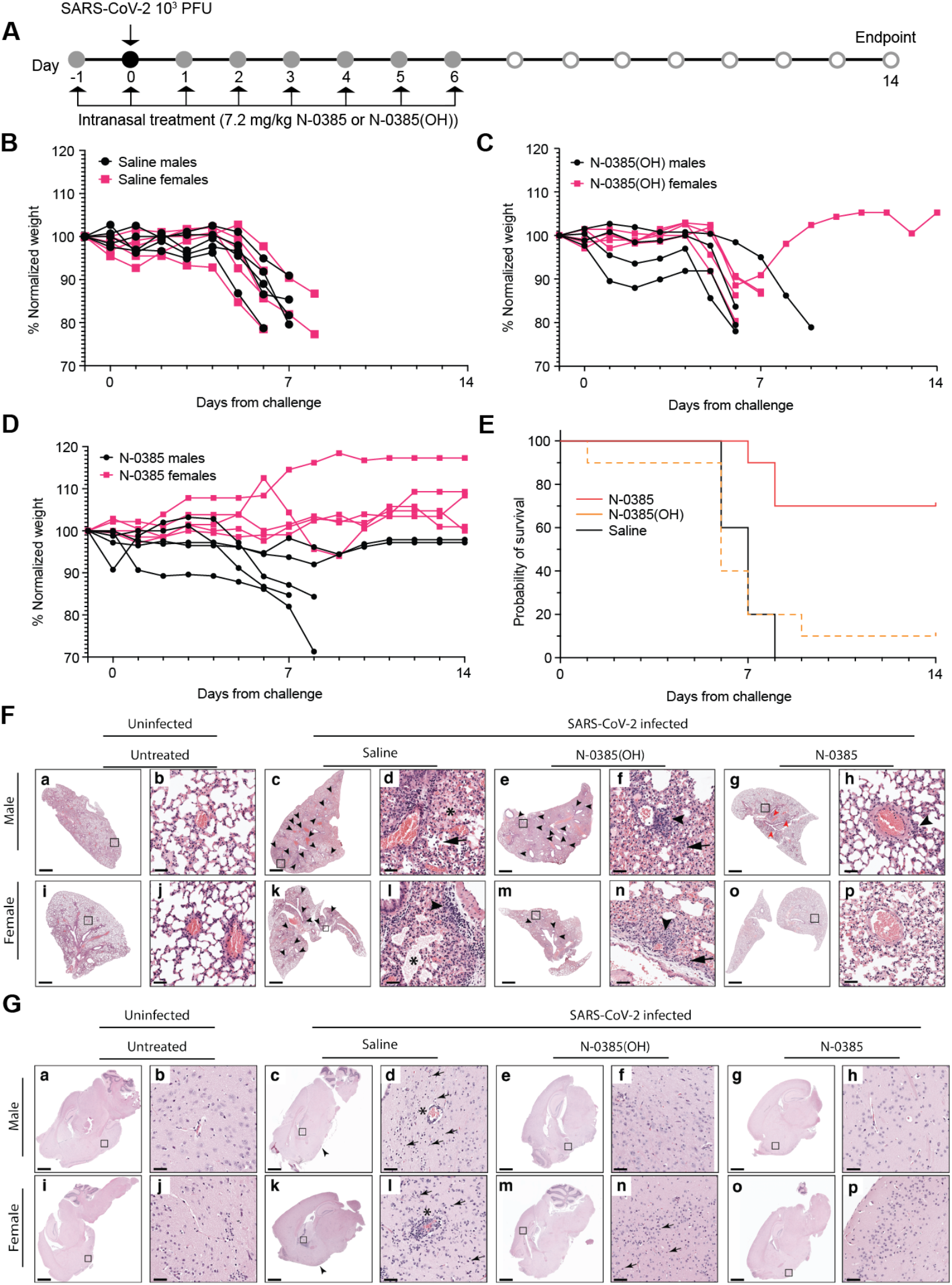
N-0385 reduces morbidity and mortality in a K18-hACE2 mice model of SARS-CoV-2 disease. **(A)** K18-hACE2 mice were treated once daily on day -1 to day 6 relative to SARS-CoV-2 infection and surviving mice were terminated on day 14. (**B)** Weight change of saline control-treated mice. (**C)** Weight change of N-0385(OH)-treated mice. (**D)** Weight change of N-0385-treated mice. (**E)** Survival graph. **(F)** Representative H&E images of lung histopathology in male (**top row**) and female (**bottom row**) mice without treatment (**a, b, i, j**) and treated with saline, N-0385(OH), or N-0385 (**c-h, k-p**). Uninfected K18 mice without treatment (**a, b, I, j**) were normal. Mice infected with SARS-CoV-2 (**c-h, k-p**) frequently developed small perivascular infiltrates of inflammatory cells (**arrowhead**). Severe inflammatory changes including alveolar fibrin and edema (**asterisk**) were present only in mice treated with saline (**c, d, k, l**). Perivascular inflammatory cell infiltrates (**arrowhead**) were more widespread in mice treated with saline (**c, k**) and N-0385(OH) (**e, m**) compared to mice treated with N-0385 (**g, o**). Mice treated with N-0385 that survived to the study endpoint (**g, h, o, p**) had smaller and fewer perivascular inflammatory infiltrates (**arrowhead**) and occasional type II pneumocyte hyperplasia (**red arrow**). (**G)** Representative H&E images of brain histopathology in male (**top row**) and female (**bottom row**) mice without treatment (**a, b, i, j**) and treated with saline, N-0385(OH), or N-0385 (**c-h, k-p**). Mice treated with saline and infected with SARS-CoV-2 (**c, d, k, l**) developed perivascular cuffs of inflammatory cells (**asterisk**), necrotic neurons (**arrow**), gliosis, and meningeal infiltrates (**arrowhead**). CNS lesions were reduced in mice treated with N-0385(OH) (**e, f, m, n**) and absent in mice treated with N-0385 that survived to the study endpoint (**g, h, o, p**). Magnified areas were selected to best represent the presence of inflammatory cells and pathological changes. Scale bar: **a, c, e, g, i, k, m, o** = 1 mm; **b, d, f, h, j, l, n, p** = 50 µm.

Histological examination of lung tissue revealed mild pathology in the majority of SARS-CoV-2 infected mice, with mild perivascular and interstitial inflammatory infiltrates as the predominant change, irrespective of treatment group (**Table S3**). Saline-treated mice frequently had additional histological changes including alveolar edema, alveolar fibrin, and inflammatory cells within alveoli. Of the mice that survived up to the study endpoint, three had focal areas of fibrosis, type II pneumocyte hyperplasia, and occasionally lymphoid hyperplasia. However, the majority of the mice that survived showed little to no pathological signs in the lungs (**Figure 4F**). Histologic lesions in the brain included multifocal perivascular cuffs of inflammatory cells, reactive glial cells, neutrophils, and lymphocytes in the adjacent neuroparenchyma (gliosis), infiltration of the meninges with inflammatory cells, and neuronal necrosis characterized by shrunken neuron bodies with hypereosinophilic cytoplasm and pyknotic or karyorrhectic nuclei. No lesions were observed in the brains of mice that survived to the study endpoint (**Figure 4G** and **Table S3**).

Immunohistochemistry of the SARS-CoV-2 nucleocapsid protein revealed significant amounts of the viral antigen throughout the brain and lungs of infected mice treated with saline or N-0385(OH) (**Figure 5** and **Table S4**). Antigen was detected in the brain, the lung, or both tissues for most of the infected mice (**Figure 5** and **Table S4)**. The amount of antigen detected in the brain and lung was reduced significantly or completely absent in mice treated with N-0385 that survived to the study endpoint (**Figure 5** and **Table S4**).

**Figure 5.**
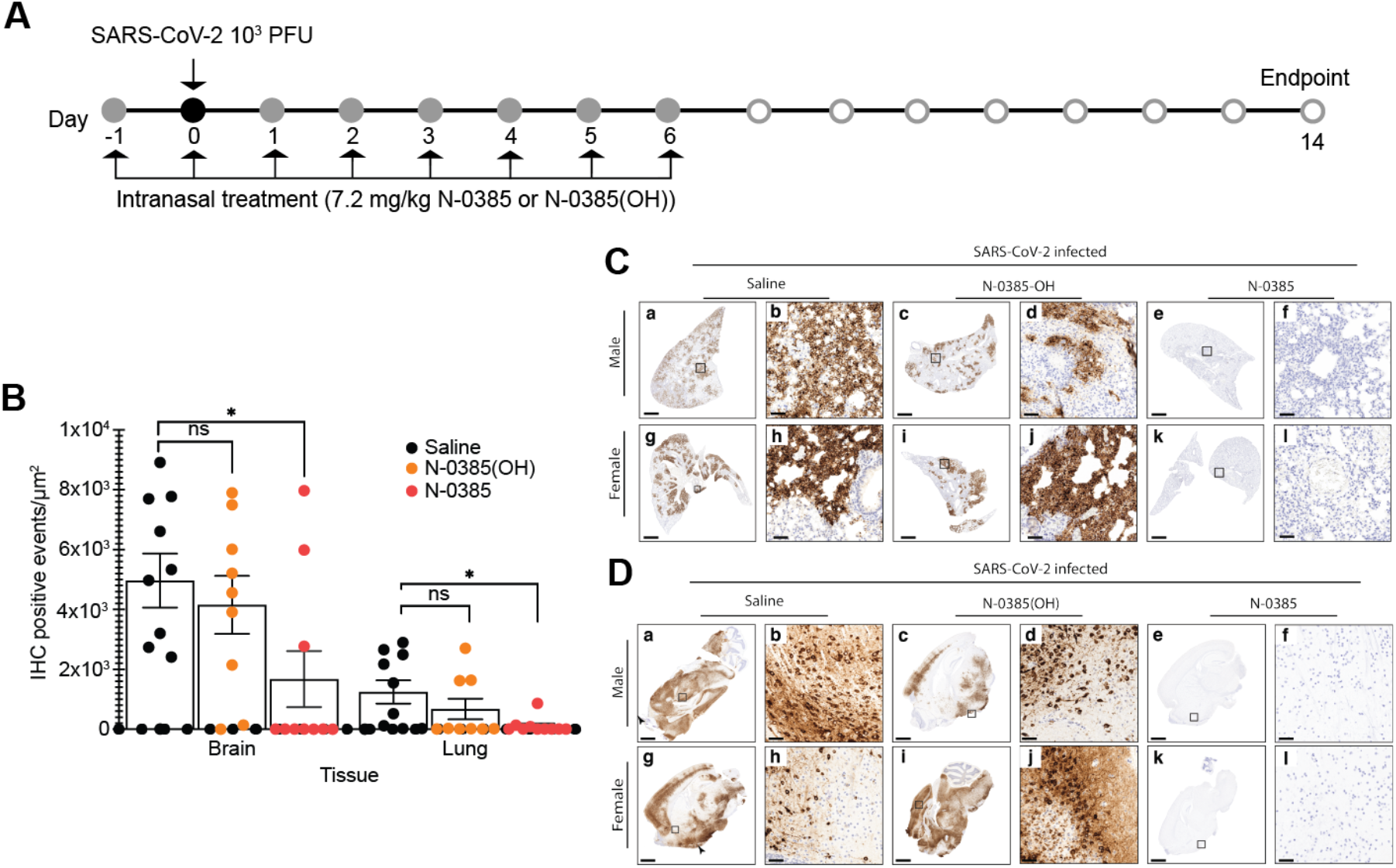
N-0385 drastically reduces SARS-CoV-2 in the lungs of mice treated with N-0385 as demonstrated by immunohistochemistry. **(A)** K18-hACE2 mice were treated once daily on day -1 to day 6 relative to SARS-CoV-2 infection and surviving mice were terminated on day 14. **(B)** Graph showing number of cells/μm^2^ that were positive for SARS-CoV-2 nucleocapsid by immunohistochemistry staining; reduction in positive cells was significantly greater for the N-0385 vs saline control while N-0385(OH) was not significantly different from the saline-treated group; * modified p <0.05. One-way ANOVA with a Bonferroni used to determine significance. Error bars indicate standard deviations. (**C**) Representative immunohistochemistry sections of SARS-CoV-2 nucleocapsid in the lungs of the SARS-CoV-2 infected male (**top row**) and female (**bottom row**) mice from Figure 3F (**a, b, g, h**). Mice treated with saline had significant immunoreactivity against SARS-CoV-2 throughout the lung. A similar pattern of patchy infection was present in mice treated with N-0385(OH) (**c, d**) but was not significant in all mice (**i, j**). Immunoreactivity for SARS-CoV-2 was rare to absent in mice that survived to the study endpoint (**e, f, k, l**). **D)** Representative immunohistochemistry sections of the SARS-CoV-2 nucleocapsid in brains of SARS-CoV-2 infected male (**top row**) and female (**bottom row**) mice from Figure 3G (**a-d, g-j**). Mice treated with saline or N-0385(OH) often had significant positive immunoreactivity in neurons throughout the brain (**e, f, k, l**). Immunoreactivity for SARS-CoV-2 was rare to absent in mice that survived to the study endpoint. SARS-CoV-2 nucleocapsid. Scale bar: **a, c, e, g, i, k** = 1 mm; **b, d, f, h, j, l** = 50 μm.

We then evaluated the outcome of a shortened regimen using ten mice per group treated with N-0385 or saline from day -1 to day 2 relative to infection (**Figure 6A**). The N-0385-treated mice in this group showed 100% survival compared to 20% survivial in the control group, while the treated group had an average 2% weight gain compared to average 14% weight loss in the control group (**Figure 6B-D**). Combined, all *in vivo* data strongly indicate that N-0385 significantly prevents morbidity and mortality in the K18-hACE2 mouse model of severe SARS-CoV-2 disease, even when treatment was limited to early infection.

**Figure 6.**
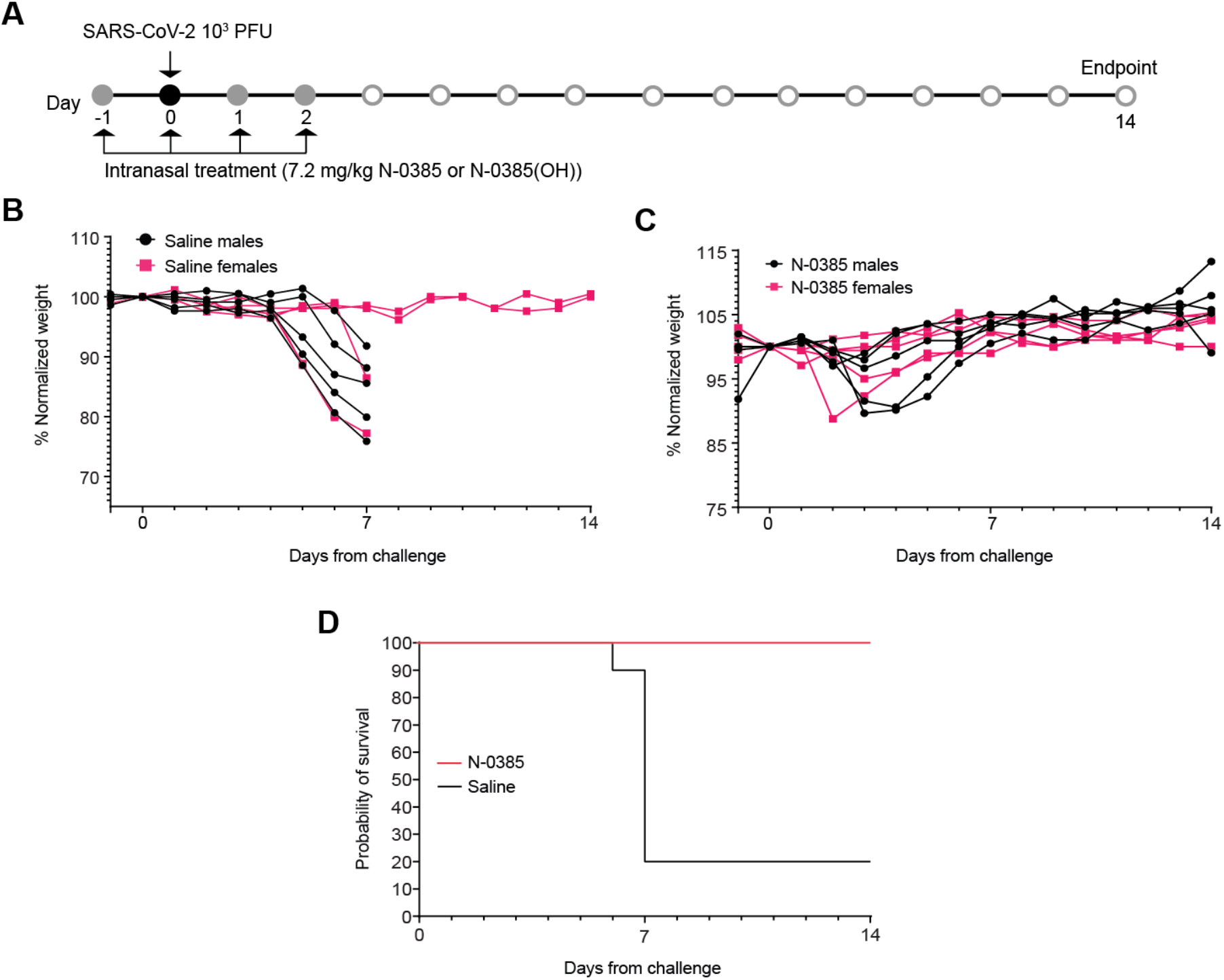
N-0385 reduces weight loss and completely prevents mortality in a K18-hACE2 mice model of SARS-CoV-2 disease following an early 4-day treatment regimen. **(A)** K18-hACE2 mice were treated once daily on day -1 to day 2 relative to SARS-CoV-2 infection and surviving mice were terminated on day 14. (**B)** Weight change of saline control-treated mice. (**C)** Weight change of N-0385-treated mice. **(D)** Survival graph. PFU = plaque-forming units

## Discussion

In the present study, we report on N-0385, the most potent small-molecule protease inhibitor of human TMPRSS2 and the first broad-spectrum nanomolar coronavirus HDA of SARS-CoV-2 VOCs. N-0385 acts as an inhibitor of the TTSP-dependent proteolytic activation of virus spike protein, a critical step to permitting viral-cell membrane fusion and entry into target cells ^4^. The nanomolar potency of N-0385 against SARS-CoV-2 infection in human Calu-3 cells and patient-derived colonoids without detectable toxicity yields a striking selectivity index of >10^6^. Furthermore, in the K18-hACE2 mouse model, treating with N-0385 resulted in complete protection against SARS-CoV-2 induced mortality, suggesting that N-0385 may provide a novel effective early treatment option against emerging SARS-CoV-2 VOCs.

We had previously shown how peptidomimetic-based compounds having ketobenzothiazole warheads exhibited potent antiviral efficacy at impeding influenza A H1N1 virus infection of Calu-3 cells through inhibition of TTSPs ^28^. The activation of the influenza A virus surface glycoprotein hemagglutinin is strikingly similar to that of the SARS-CoV-2 spike in that both are viral surface protein homotrimers cleaved by proteolytic enzymes of the TTSP family that are expressed by host epithelial cells ^14,44^. TTSPs are attractive broad-spectrum, host-directed antiviral drug targets because of (i) their important role in mediating viral entry ^5^; (ii) their accessibility on the surface of nasal and pulmonary epithelial cells ^45,46^; and (iii) their demonstrated therapeutic potential for combating medically important viruses such as SARS-CoV-2 and other human coronaviruses as well as influenza viruses ^14,44,47^.

In this work, we present the design and use of small molecule peptidomimetics with ketobenzothiazole warheads, which led to the identification of N-0385, a compound with potent inhibitory activity against TMPRSS2 proteolytic activity (IC_50_ = 1.9 nM). When we screened selected TMPRSS2 inhibitors for antiviral activity against SARS-CoV-2, a similar inhibitory profile was observed against TMPRSS2 expressed in Vero E6 cells compared to SARS-CoV-2 infection in Calu-3 cells. N-0385, the lead antiviral candidate, demonstrated potent inhibition of SARS-CoV-2 infection in Calu-3 cells, with an ED_50_ of 2.8 ± 1.4 nM and a SI of > 1 x 10^6^. The potency of this compound was validated using two viral biomarkers of intracellular infection as well as by measuring release of infectious viral particles. Further, complete inhibition of infection was achieved with 100 nM N-0385 in colonoids derived from human donors confirming the low nanomolar potency of N-0385 against SARS-CoV-2. To date, GC-376, a DAA targeting 3CL^pro^, is the only other lead antiviral candidate for SARS-CoV-2 for which a comparable SI to N-0385 has been reported in bioRxiv ^48^.

The usefulness of N-0385 needs to be envisaged in the context of the currently circulating SARS-CoV-2 variants. For example, B.1.1.7 and B.1.351 are of concern because of their rapid rise to dominance as well as their extensive spike mutations, which could lead to conformational changes of the trimeric spike structure, which may be detrimental to antiviral effectiveness and vaccine protection ^2,12,49^. We speculate that N-385 inhibitory efficacy against these two SARS-CoV-2 VOCs should not be compromised since no mutations in the TMPRSS2 cleavage site have been reported for these two SARS-CoV-2 VOCs ^12^. Our results confirmed the broad-spectrum nanomolar antiviral activity of N-0385 against SARS-CoV-2 B.1.1.7 and B.1.351 in human cells.

Recent studies have shown that the K18-hACE2 mouse model used in our studies is an ideal model to recapitulate severe human COVID-19 pathology, high morbidity and mortality. SARS-CoV-2 challenge in this model leads to high viral titers in lung and brain tissues with commensurate high morbidity and mortality and cytokine/chemokine production ^41,50^. Therefore, this model is ideal for testing SARS-CoV-2 therapeutics due to its severe disease burden, as compared to other animal models including mouse-attenuated SARS-CoV-2 in wild-type mice or wild-type SARS-CoV-2 in golden Syrian hamsters, which exhibit milder symptoms. Protection in an animal model with high levels of hACE2, such as the K18-hACE2 mouse model, is thus indicative of the high promise of anti-SARS-CoV-2 antivirals ^41^.

N-0385 significantly reduced morbidity and mortality in the K18-hACE2 mouse model of severe human COVID-19 pathology, following intranasal administration. To maximize potential antiviral efficacy, we first investigated the protective effect of an eight-day N-0385 treatment regimen, which protected 70% of the mice from SARS-CoV-2 induced mortality. Subsequently, we investigated a shortened early treatment regimen and observed 100% survival of these mice, underlining the potent antiviral efficacy of N-0385 and the importance of the TTSP-mediated proteolytic maturation of spike protein for SARS-CoV-2 infection *in vivo*. In addition to the reduced mortality, morbidity, and histological signs, immunohistochemical analysis indicated a significant reduction of SARS-CoV-2 nucleocapsid protein in the lungs and brain in the mice that survived. This is indicative of the effective reduction of virus propagation in both organ types in this animal model. While further studies are needed to understand the ideal time points for N-0385 administration to sufficiently reduce viral entry/propagation *in vivo*, N-0385 shows a greater-than-60% reduction in the proportion of SARS-CoV-2 infected cells in the lungs of infected mice, as indicated by IHC staining (day 6-8 post-infection). Our findings suggest that a single daily intranasal delivery of N-0385 can provide a novel effective early treatment option for COVID-19.

A number of lead antiviral candidates for SARS-CoV-2 infection are under investigation in clinical trials and in animal models but to date only one study on the DAA GC-376 has reported protection against lethal SARS-CoV-2 infection in the K18-hACE2 model ^23^. Plitidepsin, a newly discovered naturally occurring HDA protected against lung pathology in the K18-hACE2 model; however, the effect on mortality was not reported ^21^. Plitidepsin is a promising HDA, which targets the ubiquitously expressed elongation factor 1-alpha 1 and has demonstrated high potency (ED_50_ = 1.62 nM and SI = 40.4) against SARS-CoV-2 infection in pneumocyte-like cells ^21^. Cm and nafamostat mesylate are also HDAs targeting serine proteases, including host TTSPs, that are undergoing human trials against SARS-CoV-2; however, no significant protection against infection was observed in the adenovirus hACE2 model (Cm) or hamster model (nafamostat mesylate) of SARS-CoV-2 infection ^4,51,52^. Recently reported clinical trial data for Cm treatment of hospitalized COVID-19 patients demonstrated a lack of impact on time to recovery and incidence of death following SARS-CoV-2 infection ^19^. Antivirals will likely need to be administered during the very early phase of COVID-19 to be effective in lowering the risk of disease progression, consistent with our short early treatment regimen in K18-hACE2 mice infected with SARS-CoV-2.

Overall, we have developed and characterized N-0385, a novel highly potent inhibitor of TMPRSS2-like proteases that blocks SARS-CoV-2 VOCs and is broadly protective against infection and mortality in mice. In addition, we demonstrated that N-0385 provides a novel effective early treatment option against emerging SARS-CoV-2 VOCs. Further, N-0385 analogs may have broader applications in combating other widespread respiratory viruses that usurp TMPRSS2-related proteases for viral entry, including other established coronaviruses, influenza viruses, and other viruses that depend on TTSPs for entering host cells ^4,30,44^. We envision a practical use of N-0385 for unvaccinated individuals or those with high risk of exposure or severe disease outcome related to SARS-CoV-2 VOCs and future emerging pathogens.

## Materials and Methods

### Cell lines, antibodies, and inhibitors

Calu-3 cells (ATCC® HTB-55™) were cultivated according to ATCC recommendations. All experiments were performed in these cells below passage 6. Vero E6 cells (ATCC^®^ CRL-I1586™; used for SARS-CoV-2 plaque assay) were cultivated in MEM supplemented with 10% FBS, 1 mM sodium pyruvate, and 0.1 nM non-essential amino acids and used at passage 19-25. All cells were expanded in a T75 flask with 5% carbon dioxide at 37°C. Cell density was kept between 0.25 and 2 million cells/mL. Camostat mesylate was obtained from MilliporeSigma. The SARS-CoV-2 nucleocapsid antibody [HL344] (GTX635679) was kindly provided by Genetex; mouse anti-dsRNA antibody (J2-1904) was purchased from Scions English and Scientific Consulting ^34^; Hoechst 33258 and secondary antibodies goat anti-mouse IgG Alexa Fluor 488 (A11001) and goat anti-rabbit IgG Alexa Fluor 555 (A21428) were obtained from Invitrogen.

### Peptidomimetic compound synthesis

Preparation of the compounds using a mixed approach of solution and solid phase synthesis is described in the supplementary materials, in addition to a synthetic scheme of analogues, NMR, HRMS, UPLC-MS retention time, structure, purity, and molecular formula strings of compounds. Amino acids and coupling reagents were obtained from Chem-Impex International (USA) and used as received. All other reagents and solvents were purchased from Sigma-Aldrich (Canada) or Fisher Scientific (USA). Tetrahydrofuran (THF) was dried over sodium benzophenone ketyl; DCM over P2O5; methanol over magnesium. Celite (AW Standard Super-Cel® NF) was obtained from Sigma-Aldrich (Canada). Thin layer chromatography was carried out on glass plates covered with silica gel (250 µm) 60 F-254 (Silicycle). Flash chromatography was carried out with Silicaflash^®^ P60 (40-63 µm, Silicyle). Chlorotrityl chloride (CTC) resin was obtained from Matrix Innovation and generally used with a loading of 1.2 mmol/g. Reactions on resin were conducted in 60 mL polypropylene cartridges (obtained from Applied Separations) and Teflon stopcocks. Reactors were gently rocked on an orbital shaker at 172 rpm during solid phase chemistry. The resin was washed with the indicated solvent for 2-5 min with 10 mL solvent per gram of resin. Purity was analyzed on a Waters UPLC H-Class with UV detection PDA equipped with an Acquity UPLC CSH C18 1.7 µm 2.1 x 50 mm^2^ column. MS spectra were recorded on a Waters SQD 2 detector (electrospray) instrument with a linear gradient of 5-95% CH_3_CN and H_2_O containing 0.1% formic acid. Final products were purified to >95% purity (UPLC-UV) using a Waters Preparative LC (Sample Manager 2767 (fraction collector); Binary gradient module 2545, with two 515 HPLC pumps and a system fluidics organizer (SFO); Photodiode Array Detector 2998: column X Select CSH Prep C18 5 µm OBD 19 x 250 mm^2^ column; buffer: A: 0.1% HCOOH in H_2_O; B: 0.1% HCOOH in ACN; flow 20 mL/min). The gradient was 10-60% of acetonitrile at a flow rate of 20 mL/min. Purities of all compounds in this paper were >95% as assessed by UPLC.

### Molecular modelling

A homology model of TMPRSS2 catalytic domain was built using the structure of matriptase (PDB: 6N4T) with the “Homology Model” module of the Molecular Operating Environment (MOE) from the Chemical Computing Group. Sequence alignment of catalytic domains of matriptase with TMPRSS2 using “Align Sequences Protein BLAST” and MOE sequence alignment allowed building of a high-quality model. Ten models were created, and the final model was selected using the best score obtained by the generalized-born volume integral/weighted surface area (GBVI/WSA) scoring method ^53^. The final model was refined and minimized using the Amber10:Extended Huckel Theory (EHT) force field. After drawing the structure, all protein-ligand complexes were prepared using the Protonate 3D tool; then the partial charges were calculated and the ligands were energy-minimized. Molecules were docked in the protein-binding site with the software MOE2019.01.02. All atoms were fixed, and the ligands were allowed to be flexible. The carbon of the ketone making the reversible covalent bond with the protein was fixed at 3.0 ± 0.1 Å of the catalytic serine to constrain the position of the ketobenzothiazole group within the binding site. The guanidine of the Arg in P1 was also fixed via two key interactions in the binding site. Conformational search using LowModeMD was made with AMBER10:EHT as a molecular mechanics force field with default parameters (rejection limit: 100; RMS gradient: 0.05; conformation limit: 10000 and iteration limit: 10000). Finally, a second round of energy minimization was performed around the ligand-binding site. The low energy conformations of the inhibitor-protein complexes were analysed for their binding interactions.

### TMPRSS2 pericellular activity screening assay and IC_50_ determination

Vero E6 cells were transfected with mock (pcDNA3.1), TMPRSS2 (pcDNA3.1/TMPRSS2 Uniprot: O15393-1), or TMPRSS2-S441A (pcDNA3.1/TMPRSS2-S441A) using Lipofectamine 3000 in 12-well plates. After 24 hr transfection, cells were washed with PBS and media replaced with HCell-100 media containing 200 µM Boc-QAR-AMC and either vehicle (0.01% DMSO) or compounds at the indicated concentration for 24 hr. To measure proteolytic activity, 90 µL of cell media was transferred to a black 96-well plate, and fluorescence was measured at room temperature (excitation: 360 nm, emission: 460 nm) using a FLx800 TBE microplate reader (Bio-Tek Instruments). Proteolytic activities are presented as percentage of activity relative to vehicle-treated cells (screen at 10 nM) or in raw fluorescence units (IC_50_ curves). IC_50_ values were determined after generating a nonlinear regression analysis from a log([Compound]) versus a proteolytic activity plot using GraphPad Prism software (version 9.0.1). GraphPad Prism was used to identify and eliminate outliers (Q = 1) and assess the goodness of the fits. IC_50_ values presented are the mean ± standard deviation (SD) of at least three independent experiments.

### SARS-CoV-2 infection and treatment in Calu-3 lung epithelial cells

All infections were carried out in a Biosafety Level 3 (BSL3) facility (UBC FINDER) in accordance with the Public Health Agency of Canada and UBC FINDER regulations (UBC BSL3 Permit # B20-0105 to FJ). SARS-CoV-2 (SARS-COV-2/Canada/VIDO-01/2020) was kindly provided by Dr. Samira Mubareka (Sunnybrook, ONT, Canada). SARS-CoV-2 VOCs (B.1.1.7 and B.1.351) were kindly provided by Dr. Mel Krajden (BC Centre for Disease Control, BC, Canada). Viral stocks were made in Vero E6 cells ^54^. For experiments, passage three of the virus was used with a determined viral titer of 1.5 x 10^7^ plaque forming units (PFU)/mL. Calu-3 cells were seeded at a concentration of 10,000 cells/well in 96-well plates the day before infection. SARS-CoV-2 stocks were diluted in cell-specific media to a multiplicity of infection (MOI) of 2. Cells were pretreated with compounds for three hr and then incubated with the virus for two days, followed by fixation of the cells with 3.7% formalin for 30 min to inactivate the virus. The fixative was removed, and cells were washed with PBS, permeabilized with 0.1% Triton X-100 for 5 min and blocked with 1% Bovine serum albumin (BSA) for 1 hr, followed by immunostaining with the mouse primary antibody J2 (dsRNA) and rabbit primary antibody HL344 (SARS-CoV-2 nucleocapsid) at working dilutions of 1:1000 for 1 hr at room temperature. Secondary antibodies were used at a 1:2000 dilution and included the goat anti-mouse IgG Alexa Fluor 488 and goat anti-rabbit IgG Alexa Fluor 555 with the nuclear stain Hoechst 33342 at 1 µg/mL and F-actin staining with Alexa Fluor 647 phalloidin at a 1:300 dilution for 1 hr at room temperature in the dark. After washing with PBS, plates were kept in dark at 4°C until imaging on a high content screening (HCS) platform (CellInsight CX7 HCS, Thermo Fisher Scientific) with a 10X objective, or a EVOS™ M7000 Imaging System (Thermo Fisher Scientific) with a 20X or 40X objective. Confocal imaging was performed with a Leica TCS SP8 STED 3× laser scanning confocal microscope (Leica, Wetzlar, Germany) equipped with a 100×/1.4 Oil HC PL APO CS2 STED White objective, a white light laser, HyD detectors, and operated with a Leica Application Suite X (LAS X) software.

### High-content screening of SARS-CoV-2 infection

Monitoring of the total number of cells (based on nuclei staining) and number of virus-infected cells (based on dsRNA and nucleocapsid staining) was performed using the CellInsight CX7 HCS platform (Thermo Fisher), as previously described ^55,56^. Briefly, nuclei are identified and counted using the 350/461 nm wavelength (Hoechst 33342); cell debris and other particles are removed based on a size filter tool. A region of interest (ROI, or “circle”) is then drawn around each host cell and validated against the bright field image to correspond with host cell membranes. The ROI encompasses the “spots” where dsRNA (485/521 nm wavelength) and SARS-CoV-2 nucleocapsid (549/600 nm wavelength) are localized. Finally, the software (HCS Studio Cell Analysis Software, version 4.0) identifies, counts, and measures the pixel area and intensity of the “spots” within the “circle.” The fluorescence measured within each cell (circle) is then added and quantified for each well. The total circle spot intensity of each well corresponds to intracellular virus levels (Z’ > 0.7) and is normalized to non-infected cells and to infected cells with 0.1% DMSO. Nine fields were sampled from each well. Nuclei stain (Hoechst 33342) was also used to quantify cell loss (due to cytotoxicity or loss of adherence) and to verify that the changes in viral infection did not result from a decrease in cell numbers.

### Median effective dose (ED_50_) curves

Intracellular dose response (ED_50_ values) for selected compounds against SARS-CoV-2 were determined by pretreating Calu-3 cells for three hr with serially diluted compounds (0.064, 0.32, 1.6, 8, 40, 200, and 1000 nM), followed by SARS-CoV-2 infection for 48 hr. Viral infection was detected by staining for dsRNA or nucleocapsid signal and quantified as described in Section *4*.*6*. ED_50_ experiments were repeated at least three times for each compound with three technical replicates in each experiment. Intracellular nucleocapsid levels were interpolated to negative control (0.1% DMSO, no infection) = 0, and positive control (0.1% DMSO, with infection) = 100. The GraphPad Prism 9™ (GraphPad Software, Inc.) nonlinear regression fit modeling variable slope was used to generate a dose-response curve [Y = Bottom + (Top-Bottom)/(1+10^((LogIC_50_-X)*HillSlope)], constrained to top = 100, bottom = 0.

### SARS-CoV-2 plaque assay

A total of 250,000 Vero E6 cells were seeded in complete MEM medium in 6-well plates and incubated for 24 hr at 37°C prior to infection with a 1:1000 dilution of supernatant from mock, infected, and treated and infected cells. The wells were washed once with PBS before 100 µL virus dilution was added per well in quadruplicate. Infected cells were incubated at 37°C for 1 hr, mixing gently every 15 min, then covered with 2 mL overlay medium of 2% Avicel CL-611 (DuPont Pharma Solutions) diluted 1:1 with 2x minimum essential media (Gibco). The cells were then incubated for three days. To fix the cells, 2 mL 8% formalin was added to each well for 30 min, following removal of the Avicel/formalin solution. Cells were gently washed with 1 mL tap water/well, followed by staining with 200 µL 1% crystal violet for 5 min. Crystal violet was removed, and the cells were washed three times with 1 mL tap water/well, then dried before the viral plaques were manually counted.

### Cytotoxicity assays

Calu-3 and Vero E6 cells (2500 or 10,000 cells for samples, 80-20,000 cells for standard curve) were seeded in 96-well plates. Following a 24-hr incubation at 37°C 5% CO_2_, cells were washed with D-PBS and compounds added (10 µM) for an additional 24-hr incubation. Cellular viability was assessed using Cell Titer-Glo® 2.0 Cell Viability Assay (Promega) according to the manufacturer’s instructions. The number of viable cells was extrapolated using the standard curve. Cellular viability in Vero E6 cells was expressed relative (%) to vehicle-treated cells. Data are from four independent experiments (mean ± SD).

### Protease selectivity of N-0385

Recombinant human matriptase, hepsin, and DESC1 were expressed and purified as described previously ^57,58^. Recombinant human furin, human cathepsin L (Bio-techne), and human thrombin (MilliporeSigma) were obtained from commercial sources. Dissociation constants (*K*_i_) were determined using steady-state velocities as previously reported ^27,29^. Assays were performed at room temperature in assay buffers (50 mM Tris-HCl pH 7.4; 150 mM NaCL; 500 µg/ml BSA for matriptase, hepsin, DESC1 and thrombin; 50 mM HEPES pH 7.4, 1 mM β-mercaptoethanol, 1 mM CaCl_2_, 500 µg/ml BSA for furin; 50 mM MES ph 6, 5 mM DTT, 1 mM EDTA, 0.005% Brij 35, 500 µg/ml BSA for Cathepsin L). To measure proteolytic activity, protease (0.25 to 1 nM) was added to the assay buffer containing different concentrations of compounds and a fluorogenic substrate (Boc-RVRR-AMC for furin, Z-LR-AMC for cathepsin L, and Boc-QAR-AMC for the other proteases). Activity was monitored (excitation: 360 nm; emission: 460 nm) using a FLx800 TBE microplate reader (Bio-Tek Instruments). If substantial inhibition occurred using a ratio I/E ≤ 10 plots of enzyme velocity as a function of inhibitor, concentrations were fitted by nonlinear regression analysis to the Morrison equation for tight-binding inhibitors. If inhibition occurred only at I/E > 10, plots of enzyme velocity as a function of substrate concentration at several inhibitor concentrations were fitted by nonlinear regression to equations describing different models of reversible inhibition (competitive, uncompetitive, non-competitive, and mixed model). The preferred model was used for *K*_i_ determination. *K*_i_ was calculated from at least three independent experiments (mean ± SD). The maximum concentration of compounds used for the assays was 10 µM.

### SARS-CoV-2 infection in human biopsy-derived colonoid monolayers

Intestinal biopsy-derived colonoids from healthy donors were obtained from the Johns Hopkins Conte Digestive Disease Basic and Translational Research Core Center (NIH NIDDK P30-DK089502) and grown according to Staab *et al*.^37^. Briefly, human colonoid monolayers were generated by combining the colonoids from one Matrigel dome (∼100 or more colonoids in a 25 µL dome). Domes were dislodged with a cell scraper in 1 mL of Cultrex Organoid Harvesting solution (Bio-techne, R&D Systems brand, 3700-100-01) and incubated for 1 hr at 4°C on a shaker at 250 rpm. After incubation, cells were diluted with an equal volume of complete media without growth factors CMGF (Advanced DMEM/F-12 (Gibco brand, Thermo Fisher Scientific 123634010), 10 mM HEPES (Invitrogen 15630-080), GlutaMAX (Gibco brand, 35050-061), and 100 U/mL of penicillin-streptomycin (Gibco brand, 15140-122)), and then centrifuged at 400 x g for 10 min at 4°C. Cells were resuspended in 50 µL/well of TrypLE Express (Invitrogen, 12604021) and then incubated for 1 min at 37°C. Following incubation, 10 mL of cold CMGF was added and the cells were pelleted by centrifugation as above and then resuspended in 100 µL per well of monolayer media (IntestiCult™ Organoid Growth Medium (Human) 06010), 10 µM of Rho Kinase inhibitor, Y-27632 (Stemcell 72304), and 50 µg/mL of gentamicin (Gibco brand, Thermo Fisher Scientific, 1510064)), and then seeded at a 1:4 dome-to-well ratio in a 96-well plate coated in 100 µL of 34 µg/mL human collagen IV (Sigma C5533). Cells were fed every two days and were used for experiments after they were fully confluent (4-5 days). Cells were treated with compounds for three hr prior to SARS-CoV-2 infection (MOI ≈ 1) for 72 hrs and then were fixed and stained for nucleocapsid and dsRNA as described in Sections *4*.*5* and *4*.*6*. Quantification was performed as described in Section *4*.*7*. Imaging was performed on the EVOS M7000 microscope using the following channels: 357/447nm for nuclear staining (Hoechst 33342), 470/525nm for dsRNA (Alexa Fluor 488), and 531/593nm for nucleocapsid (Alexa Fluor 555).

### SARS-CoV-2 infection and treatment in mice

Animal studies were carried out in accordance with the recommendations in the *Guide for the Care and Use of Laboratory Animals* of the National Institutes of Health. All protocols were performed under approved BSL-3 conditions and approved by the Institutional Animal Care and Use Committee at Cornell University (IACUC mouse protocol # 2017-0108 and BSL3 IBC # MUA-16371-1). Intranasal virus and antiviral treatments were performed under anesthesia, and all efforts were made to minimize animal suffering. Eight-week-old heterozygous K18-hACE2 c57BL/6J mice (strain: 2B6.Cg-Tg(K18-hACE2)2Prlmn/J) ^42,59,60^ were used for this study (Jackson Laboratory, Bar Harbor, ME). Mice were intranasally inoculated with 1×10^3^ PFU/animal using passage 1 of virus propagated in Vero E6 cells from isolate USA-WA1/2020 (BEI resources; NR-52281). Mice were housed five per cage and fed a standard chow diet. Daily treatments were administered intranasally at 7.2 mg/kg using the average weights of each group separated by sex from 1) day -1 to day 6 relative to infection (total of eight treatments) or 2) from day -1 to day 2 relative to infection (total of four treatments). Mice were monitored and weighed daily and euthanized at predetermined criteria for humane euthanasia following approved protocols, generally when weight loss reached 20% from day of challenge or mice became moribund with a clinical score >3 on a 5-point scale ^61^.

### Mice histopathology

For histologic examination, mouse lungs and brains were collected directly after euthanasia and placed in 10% formalin for >72 hrs after which tissues were embedded in paraffin. Tissue sections (4 μm) were analyzed after staining with H&E and scored blinded by an anatomic pathologist. For lung, scores were applied based on the percentage of each tissue type (alveolus, vessels, etc.) affected using the following criteria: (0) normal; (1) <10% affected; (2) 10-25% affected; (3) 26-50% affected; and (4) >50% affected. For brains, histologic scoring was assessed for perivascular inflammation using the most severely affected vessel and the following criteria: (0) no perivascular inflammation; (1) incomplete cuff one cell layer thick; (2) complete cuff one cell layer thick; (3) complete cuff two to three cells thick; and (4) complete cuff four or more cells thick. Necrotic cells in the neuroparenchyma were assessed per 0.237 mm^2^ field using the most severely affected area and the following criteria: (0) no necrotic cells; (1) rare individual necrotic cells; (2) fewer than 10 necrotic cells; (3) 11 to 25 necrotic cells; (4) 26 to 50 necrotic cells; and (5) greater than 50 cells.

To detect viral antigen, sections were labeled with anti-SARS-CoV-2 nucleocapsid protein rabbit IgG monoclonal antibody (GeneTex; GTX635679) at 1:5000 dilution and processed using a Leica Bond Max automated IHC stainer. Leica Bond Polymer Refine Detection (Leica; DS9800) with DAB was used as the chromogen. Image acquisition was performed using a Roche Ventana DP200 slide scanner. Digital image analysis was performed using QuPath software version 0.2.3 ^62,63^. Tissues were annotated to include all available lung tissue or all brain tissue excluding cerebellum, as cerebellar tissue was not available for all mice. Following annotation, automated detection was performed using automated SLIC superpixel segmentation with a DAB mean detection threshold of 0.18892.

## Acknowledgments

The authors acknowledge the support of the CL-3 facility (Facility for Infectious Disease and Epidemic Research (FINDER) of the Life Sciences Institute of the University of British Columbia founded by Dr. François Jean and its biosafety support staff including Dr. Bintou Ahidjo (Research Platform Manager) and T. Dean Airey (FINDER Senior Research Technician). We thank the LSI imaging facility of the Life Sciences Institute of the University of British Columbia funded by the Canadian Foundation of Innovation and BC Knowledge Development Fund as well as a Strategic Investment Fund (Faculty of Medicine, University of British Columbia). We further thank Dr. Alex Ball, Jr., MD, Senior Scientist (Genetex), for supplying the SARS-CoV-2 (COVID-19) nucleocapsid antibody [HL344] (GTX635679). We also thank Dr. Samira Mubareka, (Sunnybrook Health Sciences Centre and Research Institute, University of Toronto) for providing the SARS-CoV-2 (SARS-COV-2/Canada/VIDO-01/2020) and Dr. Martin Petric, Samantha Kaweski, Dr. Paul Levett, and Dr. Mel Krajden (BC Centre for Disease Control, BC, Canada) for isolating and providing the SARS-CoV-2 VOCs (B.1.1.7 and B.1.351). We also acknowledge a generous donation towards the purchase of the Cell-Insight CX7 HCS system provided by the Vancouver General Hospital Foundation to Dr. Jean. The authors also acknowledge the support of the Cornell BSL-3, animal, and biosafety support staff and the Avery August lab for their tremendous support in setting up the animal studies, including but not limited to Julie Sahler, Amie L. Redko, Nicole Kushner, Paul Jennette, Joshua Turse, Bhupinder Singh, Donna Miller, David G. Collins, Timothy Lynn Van Deusen. The authors acknowledge the support of NEOMED/adMare in the early phases of developing host-based antivirals. We also acknowledge Marco Straus and Derek Wei for help with screening pseudoparticles. We also acknowledge Eloic Colombo and Baptiste Plancq for the synthesis of some intermediates, analogs, and warhead. We also thank Dr. Jill Kelly for proofreading of the manuscript. We thank Dr. Nicholas Zachos from John Hopkins University for kindly providing human colonoids and also to the patients who donated these samples for research. The authors dedicate this work to the memory of Professor Eric Marsault ^64^ (DOI: 10.1021/acs.jmedchem.1c00481).

## Author contributions

Conceptualization: FJ, RL, HCA, PLB, GRW, EM

Methodology: TS, SD, IAM, MJ, SPD, AD, TV, DB, BI, GG, AC, PLB, JS

Investigation: TS, SD, IAM, MJ, SPD, AD, TV, DB, BI, PLB

Visualization: TS, AO, GG, AC, PLB, IAM, MJ, PLB

Funding acquisition: FJ, RL, GRW, HCA, AA, EM, IRN

Project administration: AD, AO, PLB

Supervision: FJ, RL, HCA, PLB, GRW, EM, AA, IRN

Writing – original draft: AO, FJ, PLB, RL, AD, HCA, AC, IAM, MJ

Writing – review and editing: TS, GRW, RL, PLB, AD, HCA, AC, IAM, AA, MJ, IRN, AO, FJ

## Materials & Correspondence

Correspondence and material requests related to cell-based SARS-CoV-2 studies should be addressed to Dr. François Jean, those related to peptidomimetics and *in vitro* studies should be addressed to Dr. Richard Leduc, and those related to animal studies should be addressed to Dr. Hector C. Aguilar.

